# Impact of isolation method on cellular activation and presence of specific tendon cell subpopulations during *in vitro* culture

**DOI:** 10.1101/2021.03.07.434268

**Authors:** Anne E.C. Nichols, Sarah E. Miller, Luke J. Green, Michael S. Richards, Alayna E. Loiselle

## Abstract

Tendon injuries are common and heal poorly, due in part to a lack of understanding of fundamental tendon cell biology. A major impediment to the study of tendon cells is the absence of robust, well-characterized *in vitro* models. Unlike other tissue systems, current tendon cell models do not account for how differences in isolation methodology may affect the activation state of tendon cells or the presence of various tendon cell sub-populations. The objective of this study was to characterize how common isolation methods affect the behavior, fate, and lineage composition of tendon cell cultures. Tendon cells isolated by explant exhibited reduced proliferative capacity, decreased expression of tendon marker genes, and increased expression of genes associated with fibroblast activation compared to digested cells. Consistently, explanted cells also displayed an increased propensity to differentiate to myofibroblasts compared to digested cells. Explanted cultures from multiple different tendons were substantially enriched for the presence of scleraxis-lineage (Scx-lin+) cells compared to digested cultures, while the overall percentage of S100a4-lineage (S100a4-lin+) cells was dependent on both isolation method and tendon of origin. Neither isolation method preserved the ratios of Scx-lin+ or S100a4-lin+ to non-lineage cells seen in tendons *in vivo*. Combined, these data indicate that further refinement of *in vitro* cultures models is required in order to more accurately understand the effects of various stimuli on tendon cell behavior.

**Statement of clinical significance:** The development of informed *in vitro* tendon cell models will facilitate enhanced screening of potential therapeutic candidates to improve tendon healing.

## INTRODUCTION

Following acute injury, tendons heal by the formation of a fibrotic scar. Though the scar restores tissue continuity, it can also severely limit functionality and can predispose the tendon to further injury and degeneration (1). Despite the frequency at which these injuries occur and the associated long-term complications, there are currently no treatments available that can effectively promote improved tendon healing due in part of a lack of understanding of the specific cellular mechanisms that mediate either regenerative tenogenic processes, or the pathological fibrotic response. .

In an effort to improve tendon healing, many studies have employed *in vitro* studies to better understand tendon cell behavior and to identify potential targets for therapeutic intervention, an approach which has been successfully employed in other musculoskeletal tissues such as bone, cartilage, and muscle. In these tissues, well-characterized *in vitro* model systems have led to fundamental discoveries regarding the mechanisms regulating cell differentiation (2–5) and the important role of mechanical stimuli on tissue homeostasis (6–9). These *in vitro* findings can display a high degree of translatability and have informed *in vivo* studies that now underpin our basic understanding of these tissues. Despite the critical need for robust *in vitro* model systems, there is currently no standard for the study of tendon cells. Due to a lack of suitable tendon cell lines, the field relies on the isolation and culture of primary tendon cells; however there is still no standardized protocol for tendon cell isolation, with isolation methods (explant culture or enzymatic digest) and cells derived from different tendons assumed to be equivalent and frequently used interchangeably.

It has long been understood that once isolated, tendon cells from many different species do not retain their *in vivo* characteristics, a phenomenon referred to as “phenotypic drift” (10). This process is typically described as a loss of tenogenic markers, including scleraxis (Scx), tenomodulin, Mohawk (Mkx), and collagen type 1 (Col1), increases in the expression of injury-associated collagens including collagen type III (Col3), alterations in cell morphology away from the typical spindle-shape, and changes in proliferative capacity over time in culture (10–14). This has traditionally been viewed as a negative outcome, and investigators have been advised to use isolated tendon cells at early passages in an attempt to preserve the tendon cell phenotype seen *in vivo* (12).

Though they are referred to by many different names, the majority of the cells that reside in tendons are fibroblasts. In general, when subjected to injury or stress, tissue-resident fibroblasts lose their quiescent, homeostatic phenotype and can activate to a proliferative, secretory phenotype in a well-defined process called fibroblast activation (15); however the exact manner in which this occurs as well as the ultimate fate of activated resident fibroblasts is specific to a given tissue. In general, fibroblast activation is characterized by decreased expression of tissue-specific markers and increased expression of activation markers including periostin (Postn) (16), S100a4 (fibroblast-specific protein-1; *Fsp-1*) (17), CD248 (18), and CD106 (19). While there is evidence that tendon cells activate following acute injury *in vivo* (20–22), the degree to this process is conserved in tendon cells following isolation is unknown.

Fibroblast activation following injury can ultimately lead resident fibroblasts to differentiate to a myofibroblast phenotype (23). Myofibroblasts are contractile cells that are defined by a highly organized cytoskeleton that includes the presence of alpha smooth actin (α-SMA) containing stress fibers. While myofibroblasts are crucial for wound healing to occur, prolonged or aberrant myofibroblast activity is associated with fibrotic outcomes (24), therefore making the process of fibroblast activation to a myofibroblast phenotype an attractive target for modulating healing outcomes. Recent lineage-tracing studies have shown that some tendon cells can become α-SMA+ myofibroblasts following injury *in vivo* (20) and that the prolonged presence of myofibroblasts is associated with fibrotic tendon healing (21, 25, 26). However, the specific mechanisms governing tendon cell activation and differentiation to myofibroblasts are largely undetermined.

Though tendon cells have traditionally been seen as a homogenous population, recent studies have revealed that tendon cells exhibit remarkable heterogeneity, both within a given tendon (20, 27) and between different tendons based on their anatomical location (28). In addition to a lack of understanding of how various isolation methods affect the activation status of tendon cells, there has been no characterization of how *in vivo* heterogeneity is reflected in *in vitro* tendon cell cultures to date. Conflicting data on the tendon cell response to various stimuli may be partially explained by this lack of understanding of existing *in vitro* cultures systems. Moreover, as seen with other tissue-specific culture systems, a more thorough characterization of tendon cell behavior *in vitro* may lead to an improved understanding of tendon cell function *in vivo*. The purpose of this study was therefore twofold: 1) to investigate how different isolation methods affect tendon cell activation and fate *in vitro* and 2) to determine whether the isolation methodology affects the tendon cell populations present in *in vitro* cultures.

## METHODS

### Mice

All animal studies were approved by the University of Rochester Committee for Animal Resources. *C57BL6/J* mice (000664), *S100a4-Cre* (012641), and *ROSA-Ai9 F/F* (007909) were purchased from The Jackson Laboratory (Bar Harbor, ME). *ScxCre^ERT2^* mice were a gift from Dr. Ronen Schweitzer (Oregon Health and Science University, Portland, OR). *ScxCre^ERT2^* and *S100a4-Cre* mice were crossed to the *ROSA-Ai9 F/F* strain (*ScxCre^ERT2^;Ai9 F/-*; referred to as *Scx;Ai9*, and *S100a4Cre;Ai9 F/F*; referred to as *S100a4;Ai9* mice, respectively) as previously described (20, 26) to facilitate identification of the tendon cell lineages present following isolation. Cre+ *Scx;Ai9* mice were treated with Tamoxifen (TMX; 100mg/kg; Sigma-Aldrich, St. Louis, MO) via intraperitoneal injection for three consecutive days, five days prior to isolation. Using this TMX regimen, only cells expressing *Scx* in intact tendon, at the time of TMX labeling, are labeled red (Scx-lineage [Scx-lin+]). In *S100a4;Ai9* mice, all cells that express/have expressed S100a4 at any point are labeled red (S100a4-lin+). All isolations were performed from the tendons of 10-12 week old male and female mice.

### Tendon cell isolation

Following sacrifice, both flexor digitorum longus (FDL) tendons from three *C57BL6/J* mice (6 tendons total) were harvested. Tendons were carefully dissected from the surrounding tissue and the region from the heel to just proximal to the digit bifurcation was excised. For digested cultures, three tendons were placed into 1mL of complete Fibroblast Growth Medium-2 (FGM; #CC-3132, Lonza, Basel, Switzerland) containing 0.075% w/v collagenase type 2 (C6885, Sigma), minced, and incubated with spinning for 45 minutes at 37°C. The tissue isolate was then filtered through a 70 μm filter (#43-10070-50, Pluriselect-USA, El Cajon, CA) to remove any remaining debris, pelleted, resuspended in FGM, and plated into collagen-coated (50 μg/cm^2^ Type I Rat Tail Collagen, #354236, Corning, Bedford, MA) 35mm culture dishes. To generate explant cultures, three tendons were each cut into three pieces and placed on the bottom of collagen-coated 35mm culture dishes. Explants were allowed to adhere for 4-6 hours before being covered with FGM. Within two days, cells migrated from the explanted tendon tissue onto the bottom of the culture dish. The explant tissue was removed from the cultures at the first passaging, and monolayers derived from explanted cells were cultured as described below. Cell isolations were repeated from four different cohorts of mice for a biological n=4.

### Tendon cell culture and serial passaging

Cultures were maintained in FGM at 37 °C, 5% CO_2_, and 90% humidity with complete media exchanges every two days. At 70% confluence, cells were passaged (0.05% Trypsin EDTA, #25300–054, Gibco, Waltham, MA and 5mg/mL soybean Trypsin inhibitor, #9035-81-8, Gibco) and seeded at 4,000 cells/cm^2^ into 1× 60 mm and 1× 35 mm collagen-coated culture dishes, and 1× collagen-coated chamber slide (Lab-TekII Chamber Side 154526, Thermo Fisher Scientific, Waltham, MA). At each subsequent passage, the 60 mm plate was used both to quantify the population doubling time and to seed plates for the subsequent passage, 1× 35 mm plate was harvested for RNA isolation, and the chamber slide was fixed in 4% paraformaldehyde for 15 minutes to evaluate changes in cell morphology as well as immunohistochemical staining for Ki67 and α-SMA as described below.

### Population doubling time

At each passage, the total number of cells per 60 mm plate was quantified using a hemocytometer and used to calculate the doubling time according to the following equation:

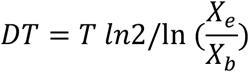

where *T* = incubation time (days), *X*_b_ = the number of cells seeded, and *X*_e_ the number of cells at passaging.

### RNA isolation and gene expression analysis

Both intact FDL tendons from six, 10-12 week old male and female *C57BL6/J* (two mice pooled per sample, n=4 samples) were snap frozen in liquid nitrogen, placed into a guanidine isothiocyanate-phenol solution (TRIzol® Reagent, Invitrogen, Carlsbad, CA) and homogenized using 0.5 mm zirconium oxide beads (# ZROB05, Next Advance Inc., Troy, NY) and a Bullet Blender Gold Cell Disrupter (Next Advance). Cultured cell monolayers from each passage were collected directly into TRIzol®. Total RNA from all samples was isolated by column purification (Direct-zol RNA Microprep kit, #R2061, Zymo Research, Irvine, CA) and converted to cDNA (qScript; #84034, Quantabio, Beverly, MA). Primers were designed (Primer Express, Applied Biosystems, Foster City, CA) validated, and used for qPCR (PerfeCTa SYBR Green; #84069, Quantabio, CFX Connect Real-Time System; Bio-Rad). Primer sequences are provided in Table 1. Fold changes were calculated using the ΔΔCt method with β*-*actin (*Actb*) as an internal control (29). Data are shown normalized to the average gene expression levels in digested cells unless otherwise stated.

**Table 1:**
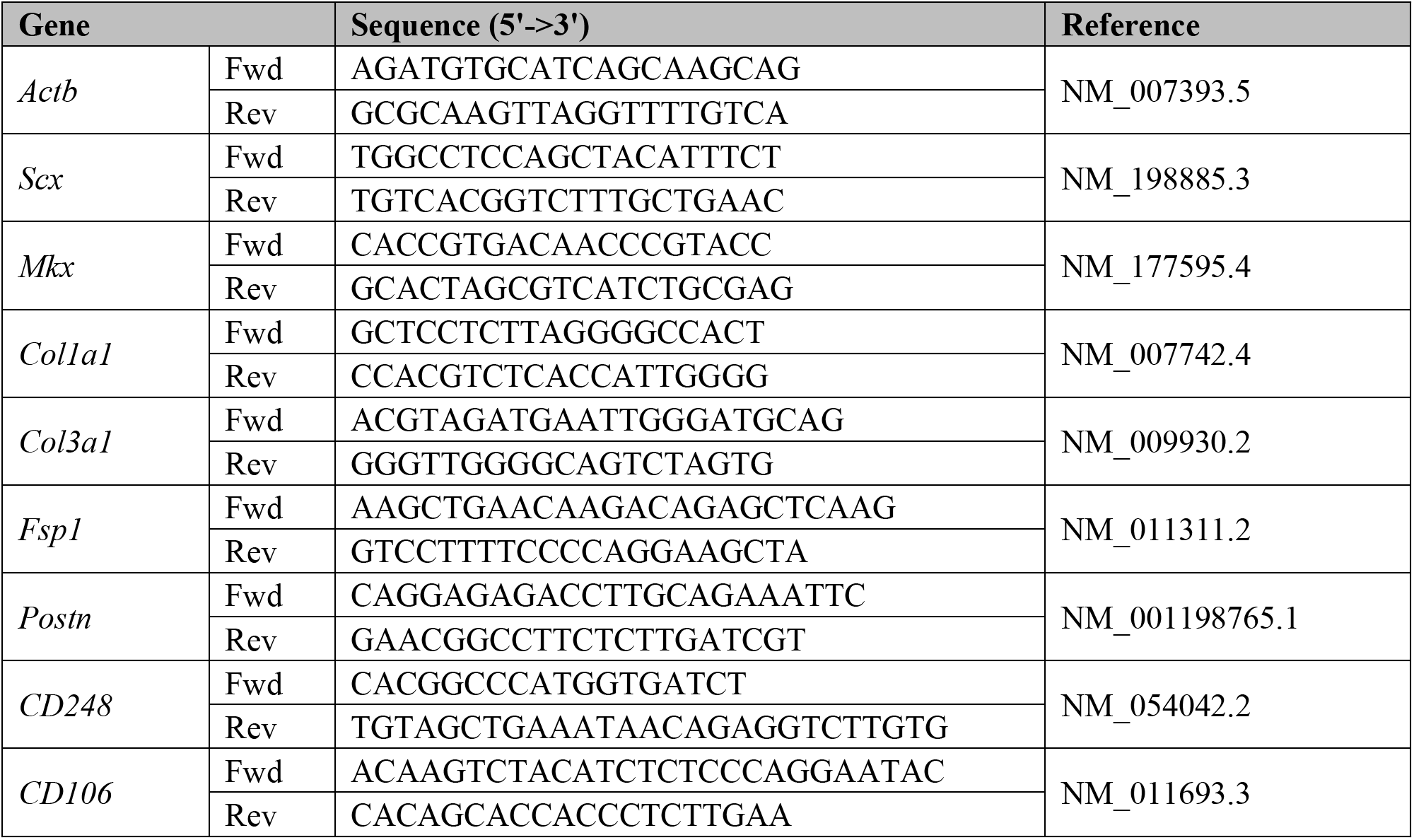
qPCR Primer Sequences

### Lineage tracing

Following sacrifice, explant and digested cultures were generated from FDL, flexor carpus ulnaris (FCU), and Achilles tendons of *Scx;Ai9* and *S100a4;Ai9* mice as described for the FDL tendon above. For tail tendons, bundles of fascicles were removed by pulling from the base of the tail tendon with sterile forceps. Explant and digested cultures were imaged every two days, and the number of lineage (red fluorescent) vs non-lineage (non-fluorescent) cells present was quantified by counting (ImageJ, National Institutes of Health, Bethesda, MD) for three random fields and averaged per culture. The lineage-tracing experiment was repeated three times (n=3).

### Immunohistochemistry (IHC)

Serially passaged cell monolayers grown on chamber slides were fixed in 4% PFA, permeabilized with 0.1% TX100, and stained with antibodies to Ki67 (1:250, ab16667, Abcam, Cambridge, MA) to evaluate cell proliferation and FITC-α-SMA (1:200, F3777, Sigma) to determine myofibroblast content. Intact front paws, hindlimbs, and tails were harvested from *Scx;Ai9* (n=4) and *S100a4;Ai9* (n=4) mice, fixed in 10% neutral buffered formalin (72 hours), decalcified in Webb Jee EDTA (14 days), processed, and embedded in paraffin. Three-micron sagittal sections were cut and exposed to primary antibodies for red fluorescent protein (1:500, AB8181-200, SICGEN, Cantanhede, Portugal) to detect the Ai9 label. Nuclei were counterstained with DAPI. Slides were scanned with a VS120 Virtual Slide Microscope and OlyVIA software (Olympus Life Science, Shinjuku, Tokyo, Japan). The number of lineage (red and blue colocalization) vs. non-lineage (blue only) nuclei within the tendon tissue were quantified using Visiopharm (Visiopharm Corporation, Westminster, CO). To confirm the localization of lineage cells within the tendon tissue, following IHC for the Ai9 label, the coverslips were carefully removed from the slides and sections were with stained with Masson’s Trichrome. For *in vitro* lineage-tracing studies, the number of Ki67+ and α-SMA+ cells present in cell monolayers were quantified using a custom MATLAB script (code available upon request; MathWorks, Natick, MA).

### Statistical analysis

All statistical analyses were performed in Prism 8 (GraphPad Software, LLC., San Diego, CA). Changes in gene expression, α-SMA+ myofibroblasts, and number of Ki67+ cells between isolation methods were evaluated by unpaired t test at each timepoint. Gene expression changes between isolated cells and intact FDL tendon were analyzed by the Kruskal-Wallis and Dunn’s multiple comparisons tests. All data are shown as the mean ± standard deviation. Statistical significance was set at p ≤ 0.05.

## RESULTS

### Tendon cell morphology and proliferative capacity differ between isolation methods

Tendon cells isolated by digest exhibited a flat, spread morphology with a clearly defined nucleus (Figure 1A, top). In contrast, the majority of tendon cells that migrated out of the explanted tissue were long, thin, and more cylindrically shaped (Figure 1A, bottom). In addition to changes in cell morphology, there was a substantial decrease in the proliferative capacity of the explanted cells compared to the digested cells, indicated by increased doubling time at every passage (Figure 1B) as well as a significantly reduced percentage of Ki67+ cells in explanted versus digested cultures at each passage beyond passage 1, with the exception of passage 3 (Figure 1C; p1: p= 0.384, p2: p=<0.001, p3: p=0.867, p4: p=0.004, p5: p<0.001). Tendon cells isolated by explant stopped actively proliferating at passage 4, though cultures could continue to be replated and kept in culture through at least passage 6.

**Figure 1:**
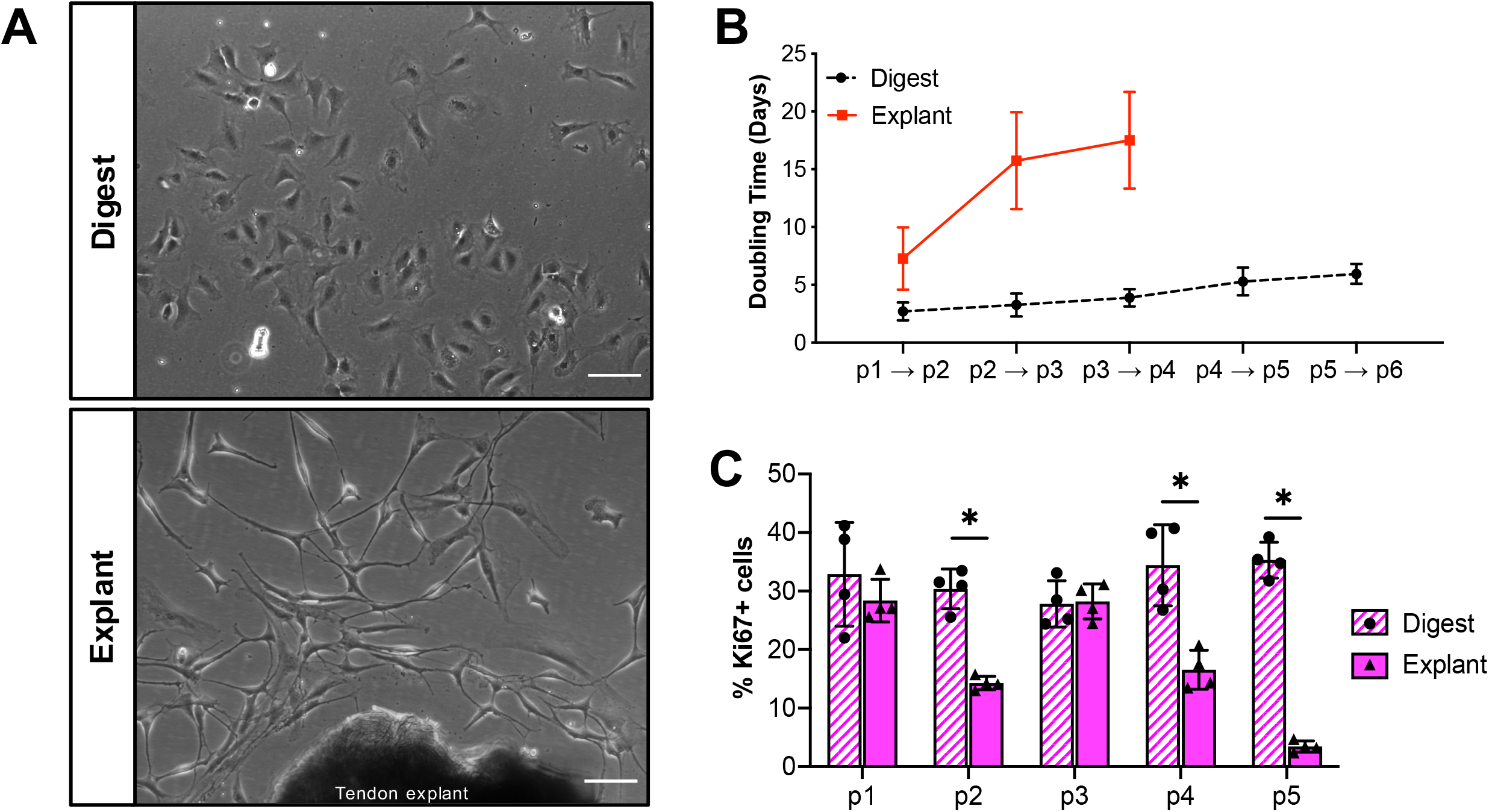
Isolation method affects tendon cell morphology and proliferation rates. (**A**) Representative images of tendon cells isolated by collagenase digest (top) or explant culture (bottom) at day 4 post-isolation. Proliferation rates of digested and explanted cells as quantified by doubling time (**B**) or Ki67+ staining (**C**) over time in culture. Scale bars = 100 μm. *p<0.05. n=4.

**Figure 2:**
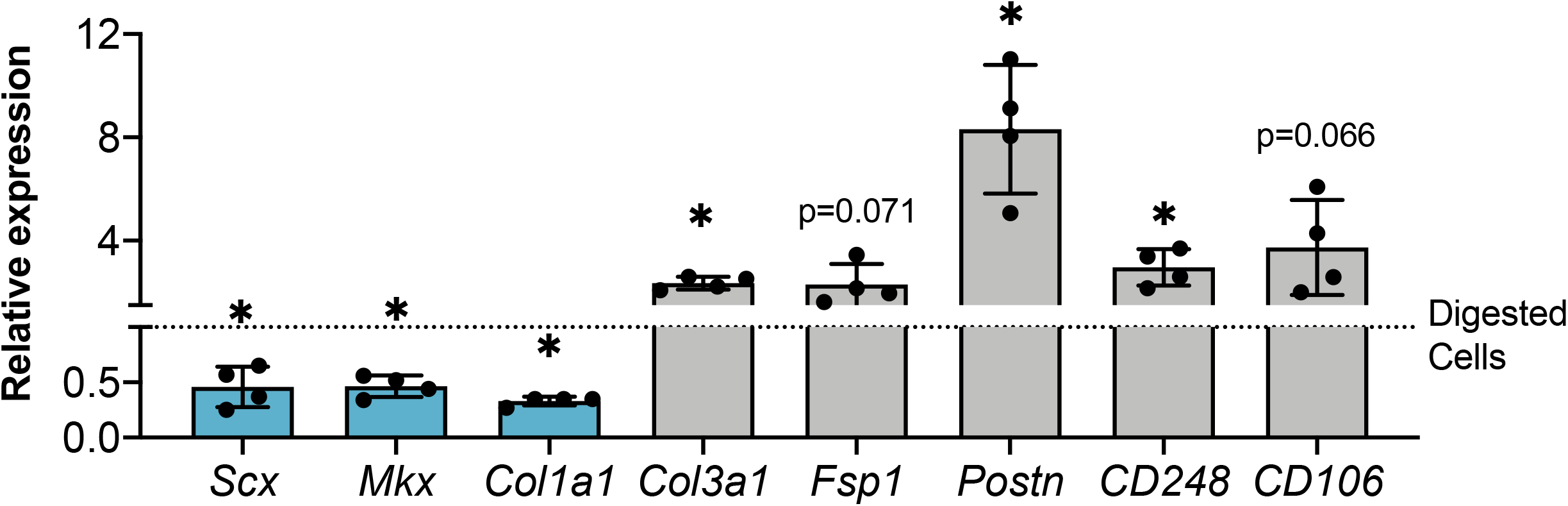
Explanted tendon cells exhibit decreased expression of tendon markers, increased expression of activation markers. Expression of tendon-related (blue) and activation marker (gray) genes in explanted tendon cells shown relative to digested tendon cells at passage 1 (dotted line). *p<0.05 compared to digested TF. n=4.

### Explanted cells exhibit a more activated gene expression profile compared to digested cells

To determine whether the changes in cell morphology and proliferative capabilities between isolation methods were reflective of changes in cell activation state, we examined the expression of tenogenic genes and genes associated with fibroblast activation at passage 1. Compared to cells isolated by digest, explanted cells exhibited significantly decreased expression of the tendon markers *Scx* (p=0.018), *Mkx* (p=0.030), and *Col1a1* (p=0.031), along with significantly increased expression of the activation markers *Col3a1* (p= 0.012), *Postn* (p=0.017) and *CD248* (p=0.015). Expression of activation markers *Fsp1* and CD106 were also increased in explanted cells compared to digested cells (~2-fold, p= 0.071 and ~4-fold, p=0.066, respectively), though these changes were not statistically significant. Digested cells also show alterations in the expression of both tenogenic markers and markers of fibroblast activation when compared to intact FDL tendon (Figure 2 supplement) suggesting that isolation by any method results in cell activation, though removal of all matrix cues via digestion results in a different activation profile than forcing cells to migrate out of explanted tissue.

### Cultures isolated by explant have a higher percentage of myofibroblasts than digested cultures

Given that one of the consequences of fibroblast activation is differentiation into a contractile myofibroblast phenotype, we next wanted to examine the effect of isolation method on myofibroblast content to determine whether the early increases in the expression of activation markers in explanted cultures relative to digested cultures would also result in increased myofibroblast differentiation. Cultures isolated by each method at each passage were stained with antibodies to the myofibroblast marker α-SMA (Figure 3A). Though the overall percentage of myofibroblasts remained low, explanted cultures contained a significantly higher percentage of α-SMA+ myofibroblasts than digested cultures at all passages examined (Figure 3B; p1: p= 0.003, p2: p=<0.001, p3: p=<0.001, p4: p<0.001, p5: p<0.001).

**Figure 3:**
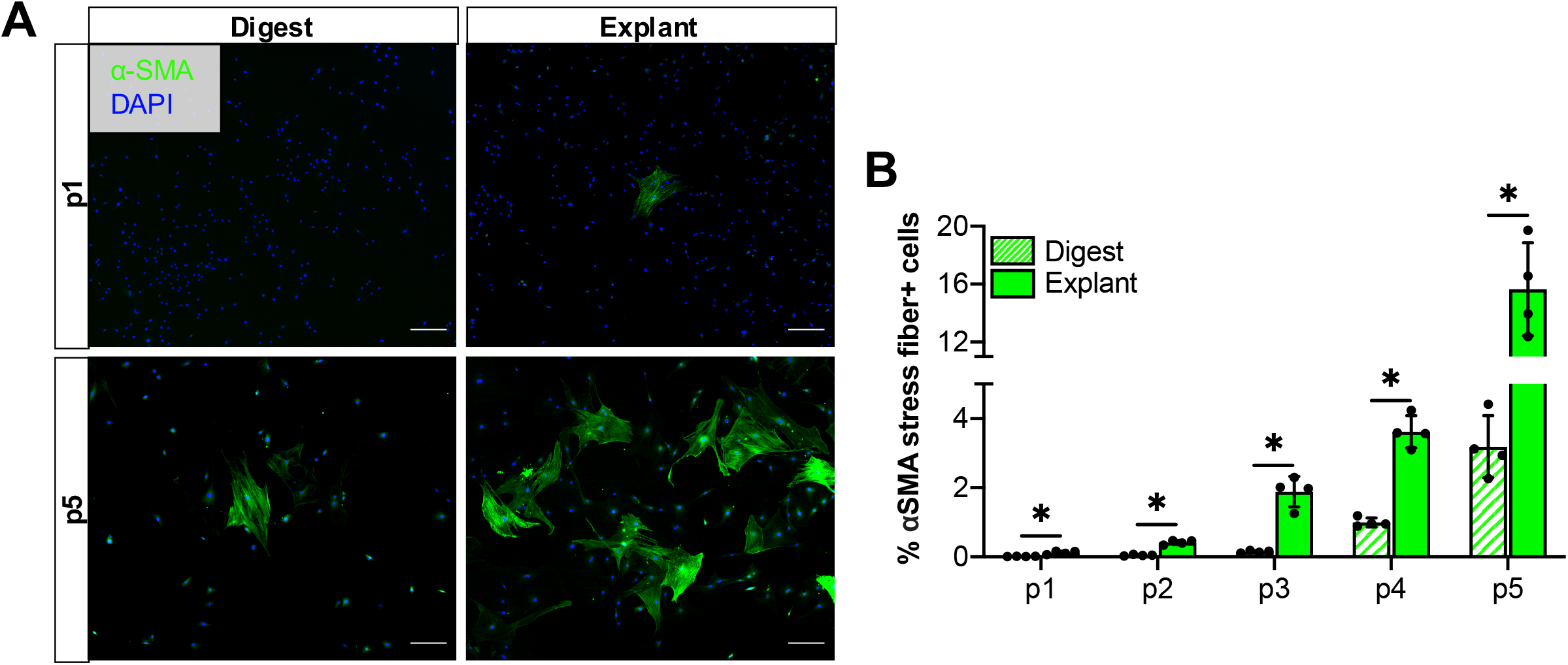
Explanted cultures contain more myofibroblasts compared to digested cultures. Representative images **(A)** and quantification **(B)** of α-SMA stress fiber (myofibroblast marker) staining over time in culture. Scale bars = 200 μm. *p<0.05, n=4.

### The composition of tendon cell lineages in vitro is dependent on isolation method

Given the recent appreciation for the heterogeneity of tendon cells (20, 27, 30), we next sought to determine if the differences in cell morphology, proliferation, activation profile and myofibroblast content between isolation methods could be due to selective isolation of different tendon cell populations. As Scx-expressing cells have traditionally been seen as the predominant resident tendon cell population, we first examined whether Scx-lineage cells made up the majority of cells in tendon cell cultures. To facilitate labeling of Scx-lineage tendon cells, *ScxCre^ERT2^+* mice were crossed to the *Rosa Ai9* reporter strain (Scx-lin+). Mice were given tamoxifen injections for three consecutive days, allowing a five day washout period prior to isolation (Figure 4A). This tamoxifen dosing regimen labeled 56.49 ± 5.14% of cells in the intact adult FDL tendon (Figure 4B). Following isolation by collagenase digest, Scx-lin+ comprised approximately 50% of the cells present through the 16 day culture period (Figure 4C). In contrast, Scx-lin+ cells made up 29.50 ± 15.49% of the cells in explanted cultures at day 2 but 93.37 ± 5.13% of cells present by day 8 post-isolation, remaining relatively unchanged from day 8 through day 16 (94.33 ± 1.55%) (Figure 4D).

**Figure 4:**
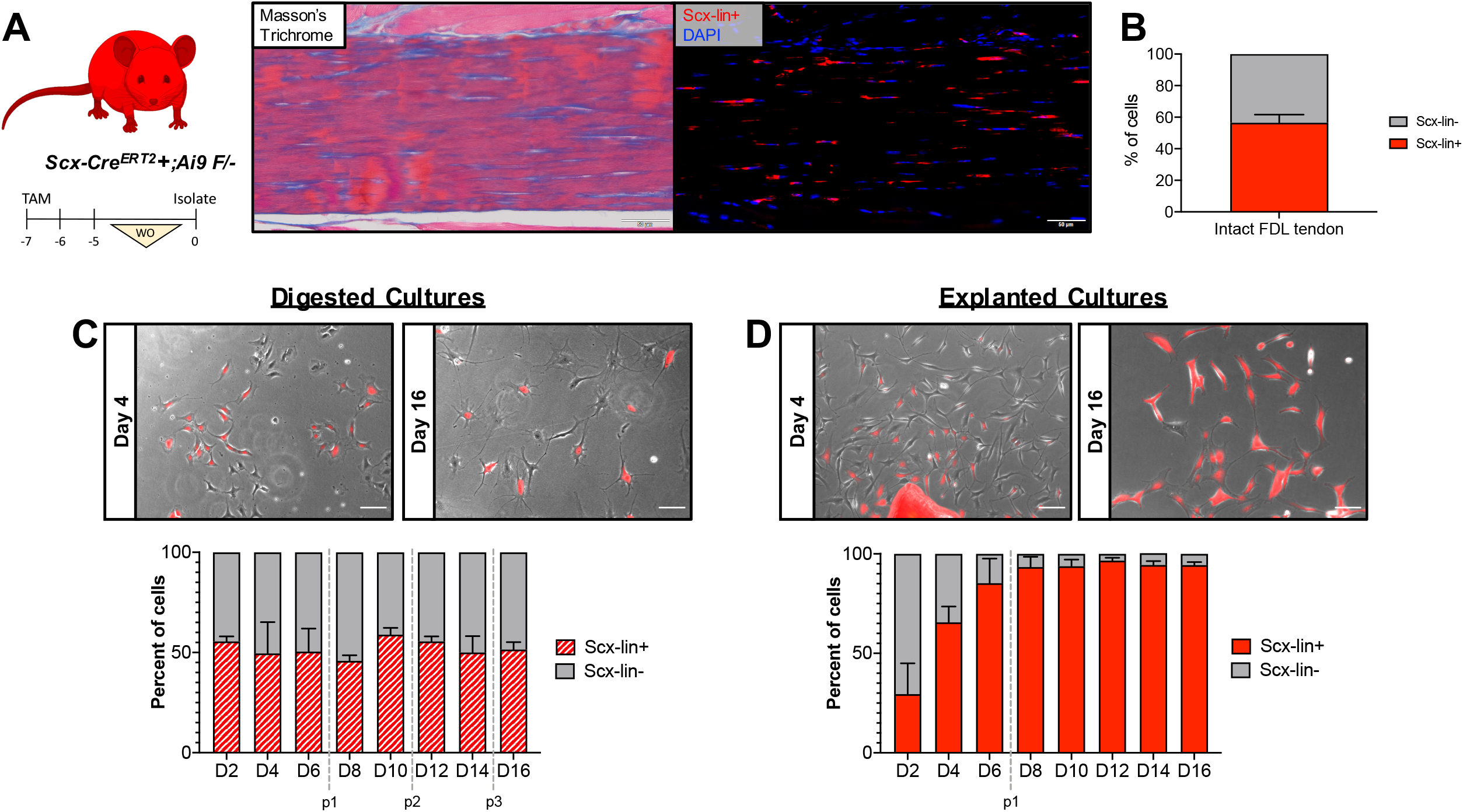
Ratio of of Scx-lin+/Scx-lin-cells in culture depends on isolation method. *Scx;Ai9* mice were injected with TMX to label Scx-lin+ cells (**A)**. Representative images showing intact FDL tendon (Masson’s Trichrome; left) and Ai9 labeling (immunofluorescence; right) of tendon cells in the same area using the described TMX regimen. Scale bars = 50 μm. Quantification (**B**) of Scx-lin+ and Scx-lin-tendon cells in intact FDL tendons. n=4. Representative images (**C,D;** top) and quantification (bottom) of Scx-lin+ (red) and Scx-lin-(gray) tendon cells isolated by digest (**C**) or explant culture (**D**). Passages are indicated by vertical gray dotted lines. Scale bars = 100 μm. n=3. Mouse model schematic was created using www.biorender.com.

Previous work has shown that S100a4 also marks a population of resident tendon cells (26), therefore we next wanted to determine the extent to which the isolation method affects the presence of S100a4-lineage cells. *S100a4-Cre* mice were crossed to the *Rosa-Ai9* reporter strain (Figure 5A) which labels 70.56 ± 6.13% of cells (S100a4-lin+) in the intact FDL tendon (Figure 5B). Similar to the Scx-lin+ cells, the percentage of S100a4-lin+ (approximately 80-90%) cells did not change over time in digested cultures (Figure 5C). In contrast, the percentage of S100a4-lin^+^ cells increased over time from 43.93 ± 8.17% at day 2 to 80.62 ± 2.37 % by day 6, remaining above 80% through day 16 post-isolation in explanted cultures (Figure 5D). Interestingly, cultures derived from both reporter strains contain a population of non-lineage cells whose presence is both isolation method and time-dependent (Figures 4C-D, 5C-D).

**Figure 5:**
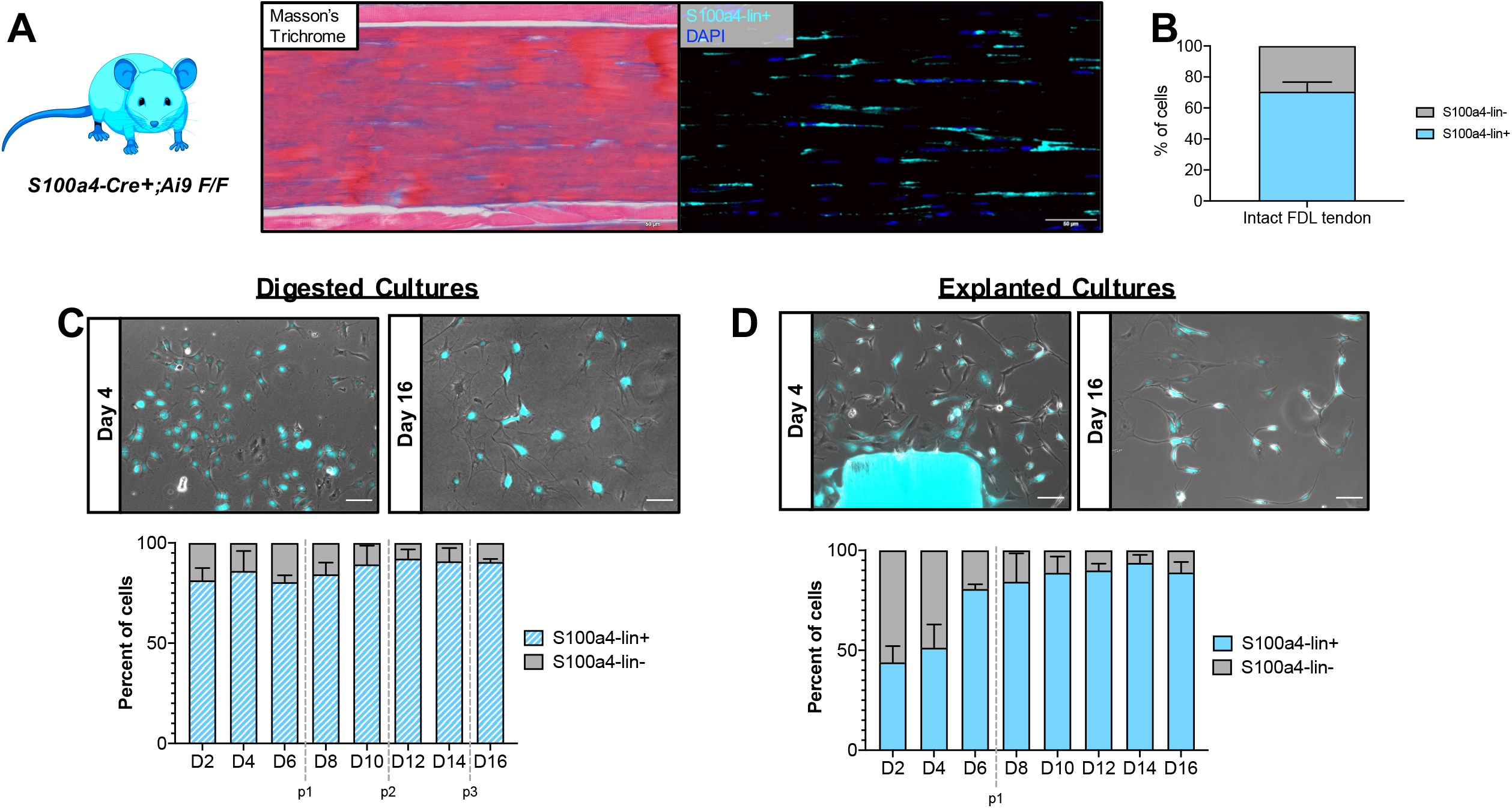
Ratio of S100a4-lin+/S100a4-lin-cells in culture depends on isolation method. Representative images (**A**) showing intact FDL tendon from *S100a4;Ai9 mice* (Masson’s Trichrome; left) and Ai9 labeling (immunofluorescence, pseudo-colored cyan; right) of tendon cells in the same area. Scale bars = 50 μm. Quantification (**B**) of S100a4-lin+ and S100a4-lin-tendon cells in intact FDL tendons. n=4. Representative images (**C,D;** top) and quantification (bottom) of S100a4-lin+ (red) and S100a4-lin-(gray) tendon cells isolated by digest (**C**) or explant culture (**D**). Passages are indicated by vertical gray dotted lines. Scale bars = 100 μm. n=3. Mouse model schematic was created using www.biorender.com.

### Scx-lin+ cell proliferation status is dependent on isolation method

Given the dramatic increase in the percentage of Scx-lin+ cells in explant cultures over time compared to the relatively steady Scx-lin+ percentage seen in digested cultures, we wanted to determine whether this difference was due to increased proliferation of Scx-lin+ cells, relative to non-lineage cells in explant cultures. Serially passaged explanted and digested cultures from *Scx;Ai9* mice were stained for the proliferation marker Ki67 and the percentage of Ki67+ cells that were Scx-lin- or Scx-lin-was quantified at each passage. Consistent with the steady proliferation seen in digested cultures over time independent of cell lineage (Figure 1C), the percentage of Ki67+ cells that were Scx-lin+ remained relatively constant, decreasing slightly over time in culture, from 86.99 ± 8.06% at passage 1 to 50.04 ± 7.84% at passage 5 (Figure 6A,B). In explanted cultures, Scx-lin+ cells made up 83.17 ± 8.40% of Ki67+ cells at passage 1, but this percentage decreased substantially over time to 19.03 ± 2.57% of cells at passage 5 (Figure 6C,D). Additionally, the percentage of Scx-lin+ Ki67+ cells was significantly lower in explanted cultures compared to digested cultures at each passage beyond passage 2 (Figure 6D, p1: p=0.535, p2: p=0.394, p3: p=0.006, p4: p=<0.001, p5: p=<0.001). Combined with the overall decrease in the proliferation of explanted cells, this similar decrease in the proliferation of explanted Scx-lin+ cells indicates that the increase in the number Scx-lin+ cells seen in explanted cultures over time is not due to on-going proliferation of Scx-lin+ cells relative to non-lineage cells.

**Figure 6:**
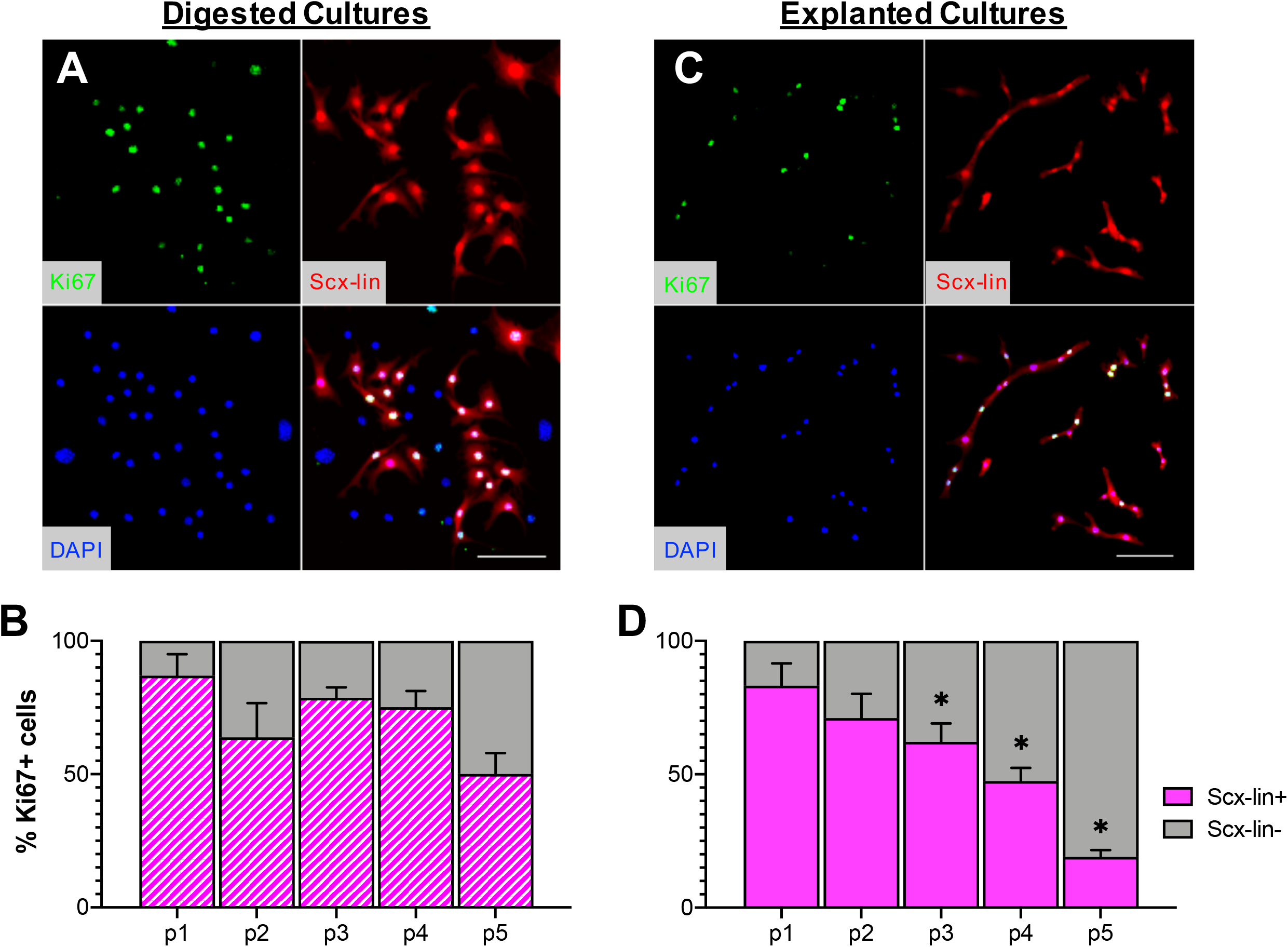
Scx-lin+ cell proliferation status is dependent on isolation method. Representative images of Scx-lin+ Ki67+ cells from digested (**A**) or explanted (**B**) cultures at passage 1. Quantification of Scx-lin+ (magenta) or Scx-lin-(gray) present over time in cultures isolated by digest (**B**) or explant **(D**) culture. Scale bars = 100 μm. *p<0.05 compared to digested cultures at the same passage. n=4.

### The majority of the myofibroblasts present in tendon cell cultures are not derived from in vivo Scx-lin+ cells

Using the same mouse model and labeling strategy employed in the current study, it has previously been reported that the majority of myofibroblasts present in the scar area of healing tendons *in vivo* are not derived from adult Scx-lin+ cells (20). To determine the Scx-lineage status of myofibroblasts present in *in vitro* cultures, explanted and digested cultures from *Scx;Ai9* mice were stained for the myofibroblast marker α-SMA to quantify the percentage of α-SMA+ myofibroblasts that were derived from Scx-lin+ cells at each passage (Figure 7A,B; shown at passage 5). Similar to what is seen *in vivo*, the majority of α-SMA+ myofibroblasts *in vitro* are not derived from Scx-lin+ cells in either digested (Figure 7C) or explanted cultures at any passage (Figure 7D); however, the percentage of Scx-lin+ myofibroblasts present did differ significantly between isolation methods. At each passage, there were significantly fewer Scx-lin+ myofibroblasts in explanted cultures compared to digested cultures (Figure 7D, p1: p=0.005 p2: 0.001, p3:<0.001, p4: 0.017, p5: <0.001).

**Figure 7:**
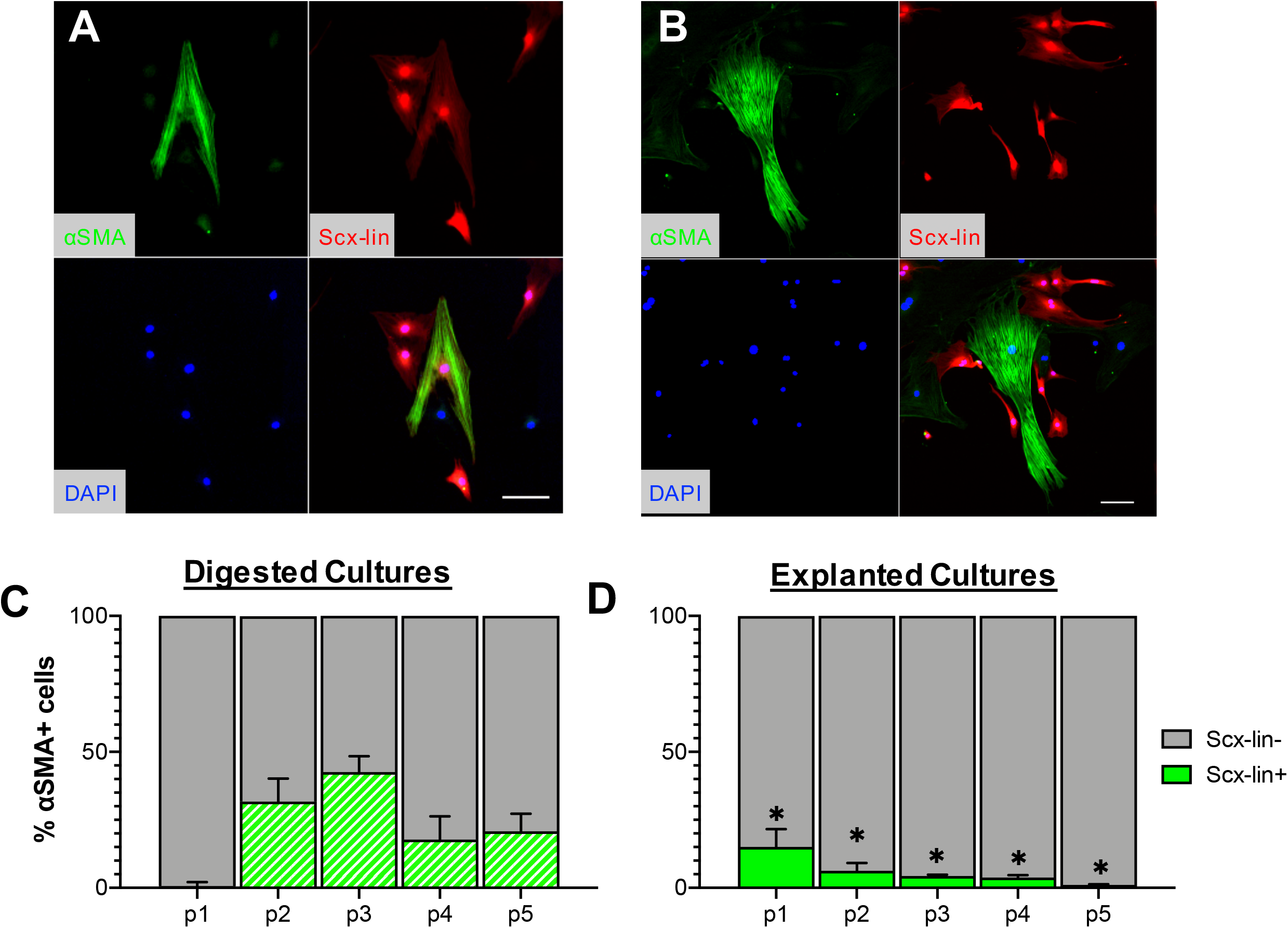
The majority of the myofibroblasts present in tendon cell cultures are not derived from *in vivo* Scx-lin+ cells. Representative images of Scx-lin+ SMA+ (**A**) and Scx-lin-SMA+(**B**) cells from passage 5 cultures. Quantification of Scx-lin+ SMA+ (green) or Scx-lin-SMA+ (gray) present over time in cultures isolated by digest (**C**) or explant **(D**) culture. Scale bars = 100 μm. *p<0.05 compared to digested cultures at the same passage. n=4.

### Tendon cell populations present in in vitro cultures are highly dependent on isolation method and the tendon of origin

Finally, we wanted to determine whether our findings regarding the effects of isolation method on tendon cell lineages present in *in vitro* cultures were specific to the FDL tendon or could also be observed across multiple tendons. We therefore performed the lineage tracing experiments with tendon cells isolated by digest or explant culture from the Achilles, flexor carpi ulnaris (FCU), and tail tendons of *Scx;Ai9* and *S100a4;Ai9* mice. To determine the *in vivo* labeling efficiency prior to isolation, sagittal sections were used to quantify the number of lin+ cells from each tendon (Figure 8). In the Achilles tendon, Scx-lin+ cells comprised 30.36 ± 5.83%, whereas S100a4-lin+ cells were 86.74 ± 1.56% of the total tendon cell population (Figure 8A-C). In the FCU, 41.55 ± 2.33% of cells were Scx-lin+ compared to 75.57 ± 4.13% that were S100a4-lin+ (Figure 8D-F). Finally, in the tail tendon, 69.40 ± 3.81% of tendon cells were Scx-lin+ with a similar 71.17 ± 2.80 cells being S100a4-lin+ (Figure 8G-I).

**Figure 8:**
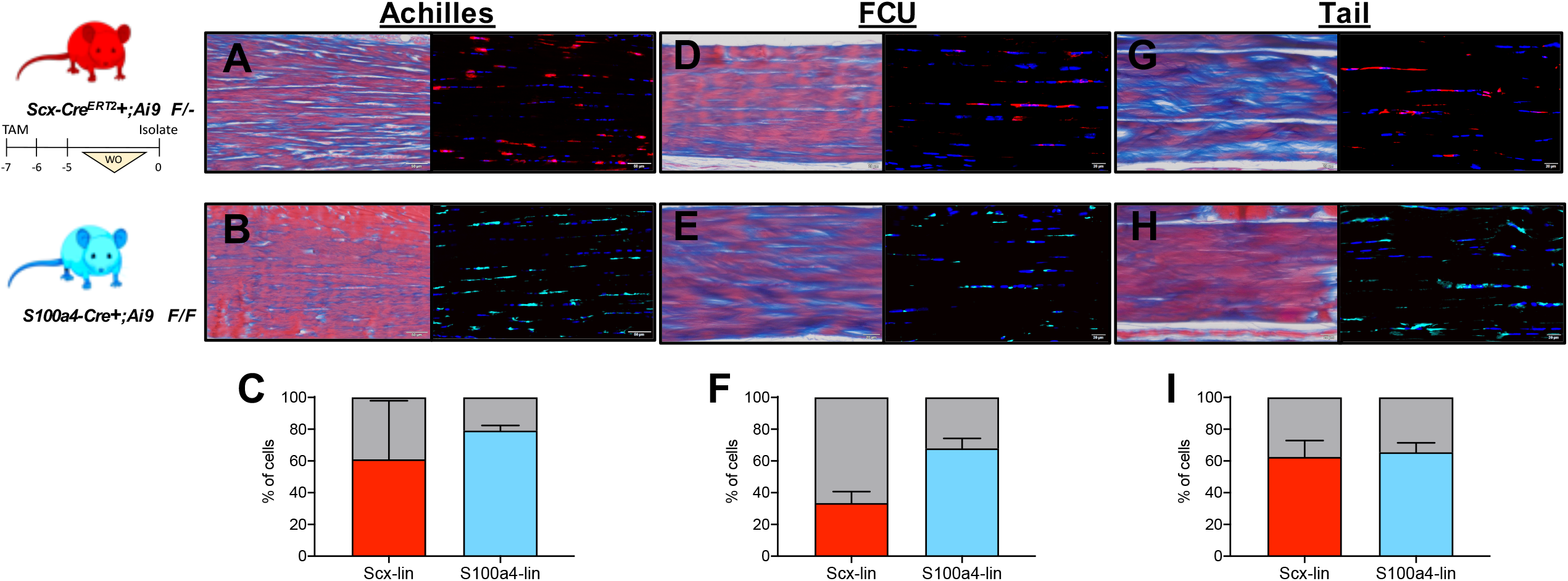
*In vivo* labeling efficiency of *Scx-Cre^ERT2^+;Ai9 F/-* and *S100a4-Cre+;Ai9 F/F* in multiple tendons. Representative images of *in vivo* labeling (left: Masson’s Trichrome, right: immunofluorescence) efficiency of *Scx-Cre^ERT2^+;Ai9 F/-* (**A, D, G**) and *S100a4-Cre+;Ai9 F/F* (**B, E, H**) in intact Achilles (**A, B**), flexor carpi ulnaris (FCU; **D, E**), and tail (**G,H**) tendons. Quantification of Scx-lin+ (red) and S100a4-lin+ (blue) in from Achilles (**C**), FCU (**F**), and tail tendons (**I**). Scale bars: A,B: 50 μm; D,E,G,H: 20 μm. Mouse model schematic was created using www.biorender.com.

Following isolation from the Achilles tendon (Figure 9A-C), a significantly higher (p<0.001) percentage of Scx-lin+ cells were present in digested (25.63 ± 2.66%) compared to explanted cultures (2.24 ± 2.67%) at day 4, but by day 16, there was a significantly higher percentage (p=0.004) of Scx-lin+ cells present in explanted cultures (58.17 ± 8.85%) compared to digested cultures (14.29 ± 9.33%). In contrast, at day 4 post-isolation from the Achilles tendon in both digested and explanted cultures, S100a4-lin+ cells comprised ~90% of the total cell population (D4 explant: 90.08 ± 1.87%, D4 digest: 92.50 ± 1.88%). By day 16 in culture, nearly all of the cells present in Achilles tendon cell cultures were S100a4-lin+ (D16 explant: 100 ± 0%, D16 digest: 96.77 ± 1.65).

**Figure 9:**
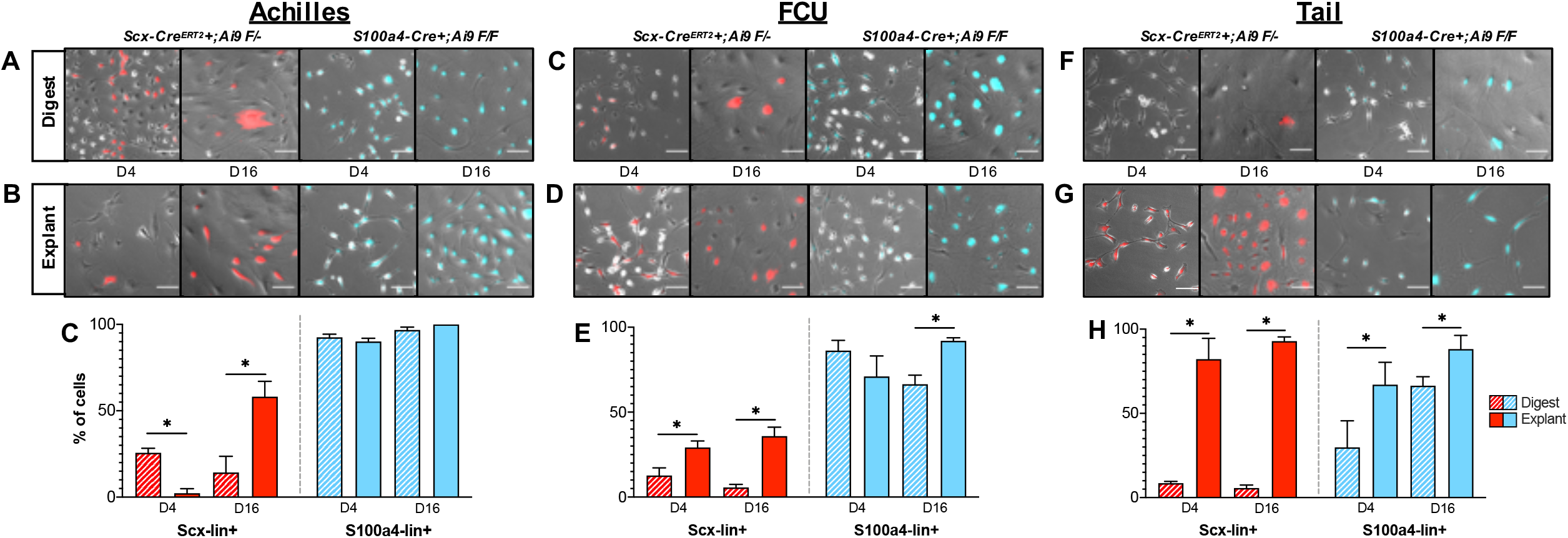
Tendon cell populations present in *in vitro* cultures are highly dependent on isolation method and the tendon of origin. Representative images of Scx-lin+ (red) and S100a4-lin+ (blue) cells isolated from murine Achilles, flexor carpi ulnaris (FCU), and tail tendons by digest (**A**) or explant (**B**) culture. Quantification of Scx-lin+ (red) and S100a4-lin+ (blue) cells by digest (striped bars) or explant culture (solid bars) at days 4 and 16 post-isolation from Achilles (**C**), FCU (**D**), and tail (**E**) tendons. *p<0.05 between isolation methods at each timepoint. n=3. Scale bar = 100 μm.

In cultures derived from FCU tendons (Figure 9C-E), explanted cultures contained a significantly higher percentage of Scx-lin+ cells compared to digested at both day 4 (p=0.008; explant: 29.20 ± 3.84%, digest: 12.67 ± 4.50%) and day 16 (p=<0.001; explant: 35.90 ± 5.25%, digest: 5.67 ± 1.80%). The percentage of S100a4lin+ cells was not significantly different (p=0.123) between explanted (70.99 ± 12.07%) and digested (86.18 ± 6.08%) at day 4 post-isolation, however at day 16, there was a significantly higher (p=0.001) percentage of S100a4-lin+ cells in explanted (91.97 ± 1.80%) compared to digested (66.43 ± 5.35%) cultures.

Following isolation from tail tendons (Figure 9F-H), explanted cultures contained a significantly higher (p<0.001) percentage of Scx-lin+ cells (82.16 ± 12.35%) than digested cultures (8.56 ± 1.05%) at day 4 which persisted through day 16 (p<0.001; explant: 92.87 ± 2.54% vs digest: 5.67 ± 1.80%). Similarly, explanted cultures contained a significantly higher proportion of S100a4-lin+ cells than digested cultures at both day 4 (p=0.035; explant: 67.07 ± 13.25, digest: 29.82 ± 15.77%) and day 16 (p=0.018; explant: 88.17 ± 8.15, digest: 66.43 ± 5.25). Combined, these data indicate that both the isolation method and tendon of origin have an enormous effect on the tendon cell populations present in *in vitro* cultures.

## DISCUSSION

To date, *in vitro* tendon cell culture systems have not taken into account how different isolations methods and source tendons may affect the ultimate phenotype and tendon cell lineage composition of cultured cells. In the present study, we examined how isolation by enzymatic digest or explant culture affect the proliferation, fate, and lineage of tendon cells isolated from the FDL tendon, demonstrating extensive differences in cellular behavior between isolation methods. Moreover, we establish that both isolation method and tendon of origin have a substantial effect on the presence of Scx-lin+ and S100a4-lin+ cells. Combined, these results suggest that further refinement of tendon cell isolation and culture protocols is required to ensure that *in vitro* studies more accurately reflect the desired tendon cell state and lineage composition.

Many previous studies have documented changes in the behavior of tendon cells following isolation, though these studies have typically been restricted to examination of tendon-specific genes. Yao *et al*. described that tendon cells isolated by collagenase digest from human Achilles tendons exhibit changes in morphology over time in culture, along with an increase in the collagen III to collagen I ratio and decreased expression of decorin (10). Cultured human tendon cells isolated by collagenase digest from the biceps long head tendon have also been shown to exhibit decreased expression of the tendon-related genes *Scx*, tenomodulin, thrombospondin-4, and Col3 along with increased expression of Col1 and tenascin-C compared to intact tendon (31). In the present study, we took inspiration from other organ-specific fibroblast culture systems and looked specifically at changes in cell behavior associated with the process of fibroblast activation. Explanted cells in our study exhibited classic hallmarks of fibroblast activation, including reduced proliferation, increased expression of activation marker genes along with decreased expression of tissue-specific genes, and increased differentiation to α-SMA+ myofibroblasts. Moreover, explanted cultures were enriched for Scx-lin+ cells in all tendons examined, suggesting that Scx-lin+ cells are the predominant migratory cell population present in tendon. Similar to what was observed in explanted cultures in this study, Best *et al.* demonstrated that adult Scx-lin+ cells migrate from the tendon ends following acute injury of the FDL tendon to form a cellular bridge and that the majority of these Scx-lin+ *in vivo* do not differentiate into myofibroblasts (20). Despite the fact that more Scx-lin+ cells were present in explanted cultures compared to digested, explanted cultures had decreased expression of *Scx*, suggesting that, as occurs following injury *in vivo* (32), activation of tendon resident Scx-lin+ cells *in vitro* involves downregulation of *Scx*. Taken together, our data suggest that explanted cell behavior *in vitro* replicates at least some aspects of the behavior of cells that migrate out of the tendon into the scar tissue *in vivo*, indicating that *in vivo* study of explanted cells could provide a valuable tool to understand tendon cell behavior following acute injury. Importantly, while explanted cells behaved similarly *in vitro* as is seen *in vivo* following acute injury, digested cells in our study also display an altered gene expression profile following isolation, though how this relates to any potential *in vivo* cell behavior is less clear. It may be that the behavior of digested cells in culture (i.e. proliferation, altered expression of activation markers relative to intact tendon) is reflective of tendon cell behavior seen in other tendinopathies *in vivo* (22), though further work is needed to confirm this. In any case, it is clear that cells isolated by these methods do not accurately represent the behavior of tendon cells during *in vivo* homeostasis.

Though previous studies have explored various isolation methods to selectively isolate different tendon cell populations based on their location within the tendon (33, 34) or their behavior *in vitro* (35–37) to our knowledge, this is the first study to utilize lineage tracing of known *in vivo* tendon cell populations to identify the tendon cell lineages present in *in vitro* cultures. Moreover, we directly compared the lineages present in cultures derived from multiple tendons isolated by both digest and explant culture. Strikingly, we observed that both isolation method and the tendon of origin have an enormous impact on the lineage of tendon cells present in *in vitro* cultures. In all tendons examined, explanted cultures were enriched for Scx-lin+ cells compared to digested cultures, with the effects of isolation methodology on the presence of S100a4-lin+ cells being dependent on the tendon of origin. Combined with the differential effects of isolation method on the activation status and fate of tendon cells *in vitro*, our lineage-tracing experiments suggest that cultures derived from different tendons by different isolation methodologies cannot, and should not, be directly compared.

Interestingly, we also identified a persistent non-lineage (lin-) cell population in cultures derived from both reporter strains whose percent of the total cell population varied depending isolation method and time post-isolation. This is particularly interesting given the differences in labeling between the two different Cre reporters: in cultures derived from the *Scx;Ai9* mice, the presence of lin- cells is consistent with the fact that not all tendon cells are Scx-lin+ *in vivo*, however in cultures derived from *S100a4;Ai9* mice, despite *Fsp-1* (also known as S100a4) being a general marker of cell activation, the presence of lin- cells (i.e. cells that have never expressed *Fsp-1* either *in vivo* or *in vitro*) suggests that *Fsp-1* is not a marker of activation in this particulcells that have never expreasr population. Future studies will clarify both the identity and role of this lin- population.

Though we have documented the effects of isolation method and tendon of origin on *in vitro* tendon cell cultures, this study is not without limitations. In all tendons examined, neither isolation method preserved the ratio of lineage to non-lineage cells seen *in vivo*, suggesting that the isolation and culture methods employed in this study select for various tendon cell populations. Though we only examined the consequence of culture in a particular culture media at a defined cellular density, changing either could provide an avenue to intentionally select for specific tendon cell lineages.

In an attempt to simulate standard tissue culture practices, we relied on passaging cells from both explanted and digested cultures at 70% confluence. Due to the decreased proliferative capacity of explanted cells relative to digested cells, this resulted in a disparity between the total time in culture between isolation methods at all passages beyond passage 1, with explanted cells at each subsequent passage taking longer to reach 70% confluence than their digested counterparts. Nonetheless, we believe that this simply highlights another issue with the lack of standardization of tendon cell culture, as passage is the most common method of reporting the ‘age’ of cell cultures, regardless of isolation method. In this study, we found stark differences between the behavior, fate, and cell lineage proportions at each passage due to isolation method, suggesting that the use of passage to report the ‘age’ of primary tendon cells in culture requires further thought and refinement in order to be able to more accurately compare the results of different studies.

In summary, the data from this study demonstrate that the both the isolation method and tendon of origin have a substantial effect on the identify and behavior of tendon cells in culture. Especially in cultures derived from explanted cells, rather than considering the changes that occur in tendon cells post-isolation as a “phenotypic drift” towards an unknown or unnatural phenotype, we propose that explanted tendon cells are in fact mimicking the natural process that occurs following injury, resulting in activation and eventual progression to a myofibroblast fate depending on the lineage from which a cell derives. The differences in activation state and the subpopulations present between isolation methods identified here may help explain discrepancies in the existing literature and should be considered when designing future *in vitro* studies of tendon cell behavior.

## ACKNOWLEDGEMENTS

The authors thank the Histology, Biochemistry and Molecular Imaging (HBMI; University of Rochester, Rochester, NY, USA) Core for technical assistance. This work was supported in part by grants from the U.S. National Institutes of Health (NIH), National Institute of Arthritis and Musculoskeletal and Skin Diseases (NIAMS) (AEL: K01AR068386 and R01AR073169, AECN: T32AR076950). The HBMI Core is supported by NIH/NIAMS Grant P30AR069655. The authors declare no competing interests.

## AUTHOR CONTRIBUTIONS

Study conceptualization and design,: AECN and AEL; data acquisition: AECN; analysis and interpretation of data: AECN, SM, LG, MSR, AEL; drafting of the manuscript: AECN; revision and approval of manuscript: AECN, SM, LG, MSR, AEL.

---

**Figure 2 (supplement):** Expression of tendon-related (blue) and activation marker (gray) genes in isolated cells at passage 1 shown relative to intact FDL tendon (dotted line). *p<0.05 between isolation methods, #p<0.05 compared to intact FDL tendon, n=4.

