## Supplementary figures and images for "Impact of isolation method on cellular activation and presence of specific tendon cell subpopulations during *in vitro* culture"

### Figure 2 supplement

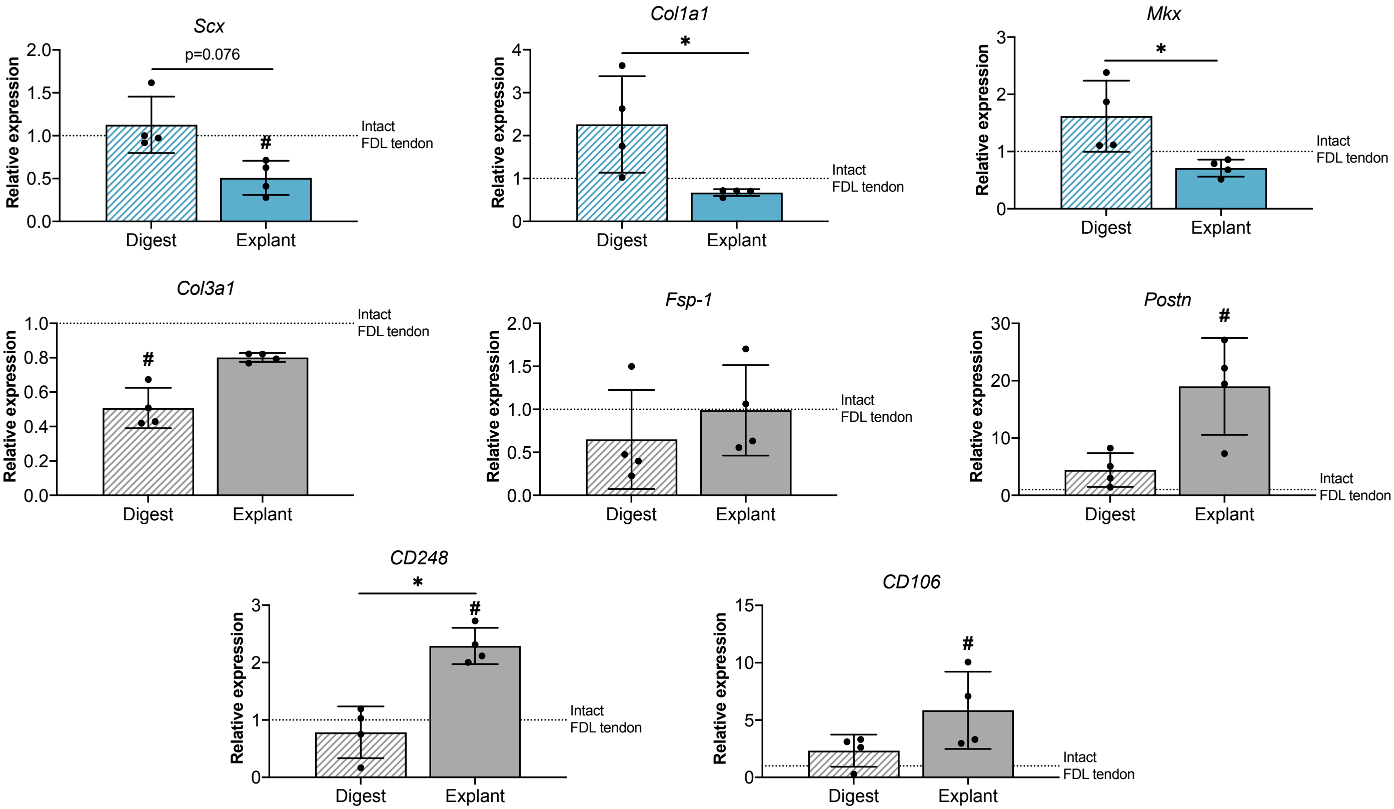
